# StereoMM: A Graph Fusion Model for Integrating Spatial Transcriptomic Data and Pathological Images

**DOI:** 10.1101/2024.05.04.592486

**Authors:** Bingying Luo, Fei Teng, Guo Tang, Weixuan Chen, Chi Qu, Xuanzhu Liu, Xin Liu, Xing Liu, Huaqiang Huang, Yu Feng, Xue Zhang, Min Jian, Mei Li, Feng Xi, Guibo Li, Sha Liao, Ao Chen, Xun Xu, Jiajun Zhang

**Affiliations:** BGI Research, Chongqing 401329, China; BGI Research, Shenzhen 518083, China; BGI Research, Hangzhou 310030, China

**Keywords:** spatial omics, multimodal data, deep learning, graph fusion, molecular characteristics

## Abstract

Spatially resolved omics technologies generating multimodal and high-throughput data lead to the urgent need for advanced analysis to allow the biological discoveries by comprehensively utilizing information from multi-omics data. The H&E image and spatial transcriptomic data indicate abundant features which are different and complementary to each other. AI algorithms can perform nonlinear analysis on these aligned or unaligned complex datasets to decode tumoral heterogeneity for detecting functional domain. However,the interpretability of AI-generated outcomes for human experts is a problem hindering application of multi-modal analysis in clinic. We presented a machine learning based toolchain called StereoMM, which is a graph fusion model that can integrate gene expression, histological images, and spatial location. StereoMM firstly performs information interaction on transcriptomic and imaging features through the attention module, guaranteeing explanations for its decision-making processes. The interactive features are input into the graph autoencoder together with the graph of spatial position, so that multimodal features are fused in a self-supervised manner. Here, StereoMM was subjected to mouse brain tissue, demonstrating its capability to discern fine tissue architecture, while highlighting its advantage in computational speed. Utilizing data from Stereo-seq of human lung adenosquamous carcinoma and 10X Visium of human breast cancer, we showed its superior performance in spatial domain recognition over competing software and its ability to reveal tumor heterogeneity. The fusion approach for imaging and gene expression data within StereoMM aids in the more accurate identification of domains, unveils critical molecular features, and elucidates the connections between different domains, thereby laying the groundwork for downstream analysis.

## INTRODUCTION

The spatial relationship between DNA/RNA and tissue-level information plays a critical role in revealing pathogenesis of cancer, developing new treat strategies, and establishing precise stratification and prognosis system. This intricate interplay allows biologists and clinicians to observe how genetic alterations manifest within the complex architecture of tissues, providing a more nuanced view of tumor biology. By integrating high-resolution genetic data with the histopathological layer, medical professionals can identify specific tumor microenvironments and spatial immune profile, as well as their responses to various treatments. This holistic approach not only aids in the development of targeted therapies that address the unique genetic makeup of the tumors but also helps in predicting disease progression and therapeutic outcomes. Consequently, leveraging the spatial dynamics between genetic information and tissue pathology paves the way for more effective and individualized cancer treatment strategies, significantly impacting patient management and improving patient survival.

The recent advent of spatial and single-cell omics technologies has produced various dimensions of information[1, 2] and indeed revolutionized our understanding of the mechanisms underpinning cancer progression and the complex tumor-immune microenvironment. These technologies provide a multi-dimensional view that captures not just the static genetic information of cells but also their spatial organization, interactions, and expression patterns within tissues (**Figure 1a**). The detailed insights provided by spatial and single-cell omics technologies into the cellular and molecular landscape of tumors represent a significant leap forward in cancer research. Spatial transcriptomics makes up for the inefficiency and accuracy of the single cell data that is resulted by the lack of in situ information. Multiple Modalities (MM) data fusion analysis paradigms have emerged in tandem with the explosion of genomics, transcriptomics, proteomics, and epigenomics. This process also has been aided by the development of artificial intelligence (AI) [3, 4]. The significant tumor heterogeneity, unpredictable drug response, and patient stratification catalyze the need for precise diagnosis and treatment of tumors, and a common trend is to combine clinical information with high-throughput data of biological and clinical level using bioinformatics and algorithms[5].

**Figure 1.**
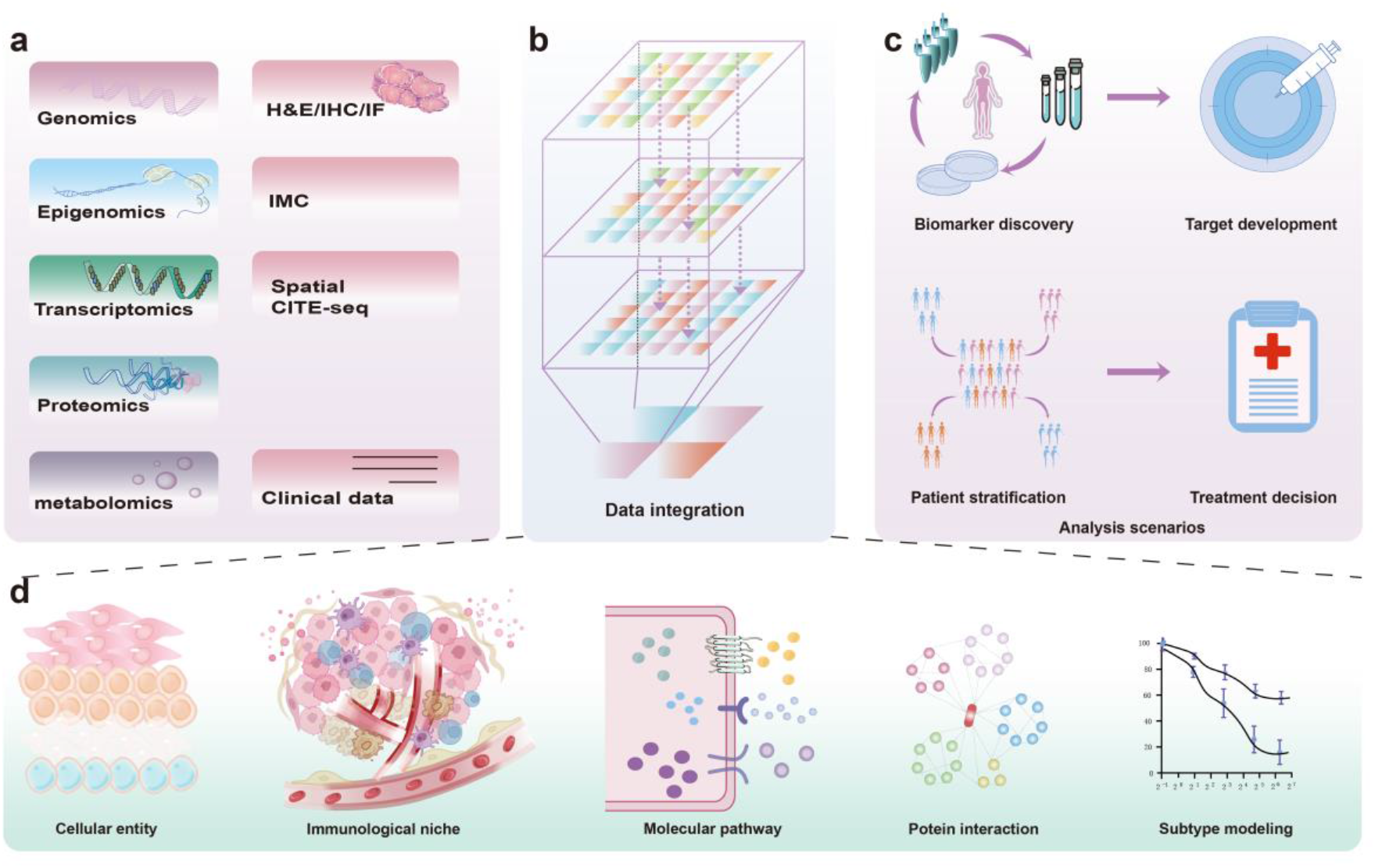
Fundamentals of Multimodal Fusion Design. **a**. Hierarchical stratification of biomedical data. **b**. Integration of aligned and non-aligned datasets. **c**. Application scenarios for multimodal data integration. **d**. Mechanistic insights via multimodal data exploration.

As spatial transcriptomic (ST) technologies develop, integration with other data modalities provide opportunities for better tissue characterization[6]. Integration of spatial transcriptomic data with conventional Hematoxylin and Eosin (H&E) histopathology images of tumor tissue opens avenues for clinical applications[7-10]. The multi-channel images provided in ST contain rich information, including cell morphology, cell status. Changes in morphology may predict cell fate or state even before it is observed in transcriptome output[11]. Meanwhile, spatial relationships between cells can reveal how different cell types and genetic programs relate to each other and their surroundings[12].

The application of multimodal data is crucial for advancing the insight into the disease and personalized cancer treatment. By integrating multiple aspects of patient information, AI algorithms can perform nonlinear analysis on these aligned or unaligned complex datasets (**Figure 1b**), achieving more precise tumor classification, disease progression prediction, and aiding physicians in crafting personalized treatment plans. This approach not only enhances the accuracy of therapeutic interventions but also facilitates the discovery of new targets and biomarkers, accelerating the development of novel drugs (**Figure 1c**). In alignment with this vision, our research endeavours extend to a granular level, where we seek to unravel the intricate biological narratives that underpin disease manifestation. We are dedicated to applications such as tumor microenvironment analysis and exploration of spatial domains, aiming to uncover the complete landscape of the disease and pioneer new avenues for treatment (**Figure 1d**).

The microenvironments specific to different regions play a pivotal role in determining cellular states, as the morphology and expression of cells reveals key insights into their physiological and phenotypic characteristics[13, 14]. Based on these assumptions, we designed the StereoMM method, which integrates RNA spatial expression data, H&E image information, and tissue in situ locations in the spatial transcriptome via cross-attention mechanisms and graph neural networks to obtain multi-modal joint embeddings. StereoMM, specifically the utilization of attention weights in the model, offers insightful explanations for its decision-making processes, thereby enhancing the interpretability of the outcomes for human experts. This feature is particularly valuable as it bridges the gap between complex algorithmic decisions and human understanding, making it possible to trace and understand the rationale behind specific predictions or classifications made by the model, thus capturing interactions between different patterns and providing a more accurate representation for downstream analysis. StereoMM has exhibited exceptional performance in identifying spatial domains. We substantiated the efficacy of StereoMM through conceptual validation across multiple cancer datasets from diverse platforms, demonstrating its superiority over existing methodologies and its potential for pivotal predictive biomarker discovery.

## RESULTS

### Overview of StereoMM framework

In the processes of diagnosis, evaluation, and therapeutic strategy formulation, physicians synthesize data from multiple sources. These data encompass three key dimensions: molecular biological, medical imaging information from clinical exams, and clinical information data from medical practice. The first dimension pertains to molecular biology, encompassing genetic, genomic, and other molecular data. The second dimension is from clinical exam, including but not limited to imaging data such as Hematoxylin and Eosin (H&E) pathology images, Immunohistochemistry (IHC), and other procedures. The last dimension is clinical practical information, which involves data derived from patient care, treatment outcomes such as response, recurrence and survival, and healthcare interactions (**Figure 1**). However, contemporary clinical diagnostic approaches may not adequately consider the potential nonlinear relationships between these different data types.

A diverse array of methodologies in spatial transcriptomics has dramatically transformed our comprehension of tissue heterogeneity and provided opportunities for multimodal fusion. Stereo-seq technology stands out for its high-resolution capabilities and expansive field of view, facilitated by a chip composed of closely spaced DNA Nanoballs (DNBs) shown in **Figure 2a**. This allows the detailed high-resolution gene expression analysis and the examination of large tissue sections, providing valuable insights into cellular heterogeneity and tissue architecture. The integration of these advantages into a multimodal data fusion algorithm framework is crucial. It merges spatially resolved gene expression data with acquired images, where structural differences could reflect functional variations, as in **Figure 2b**.

**Figure 2.**
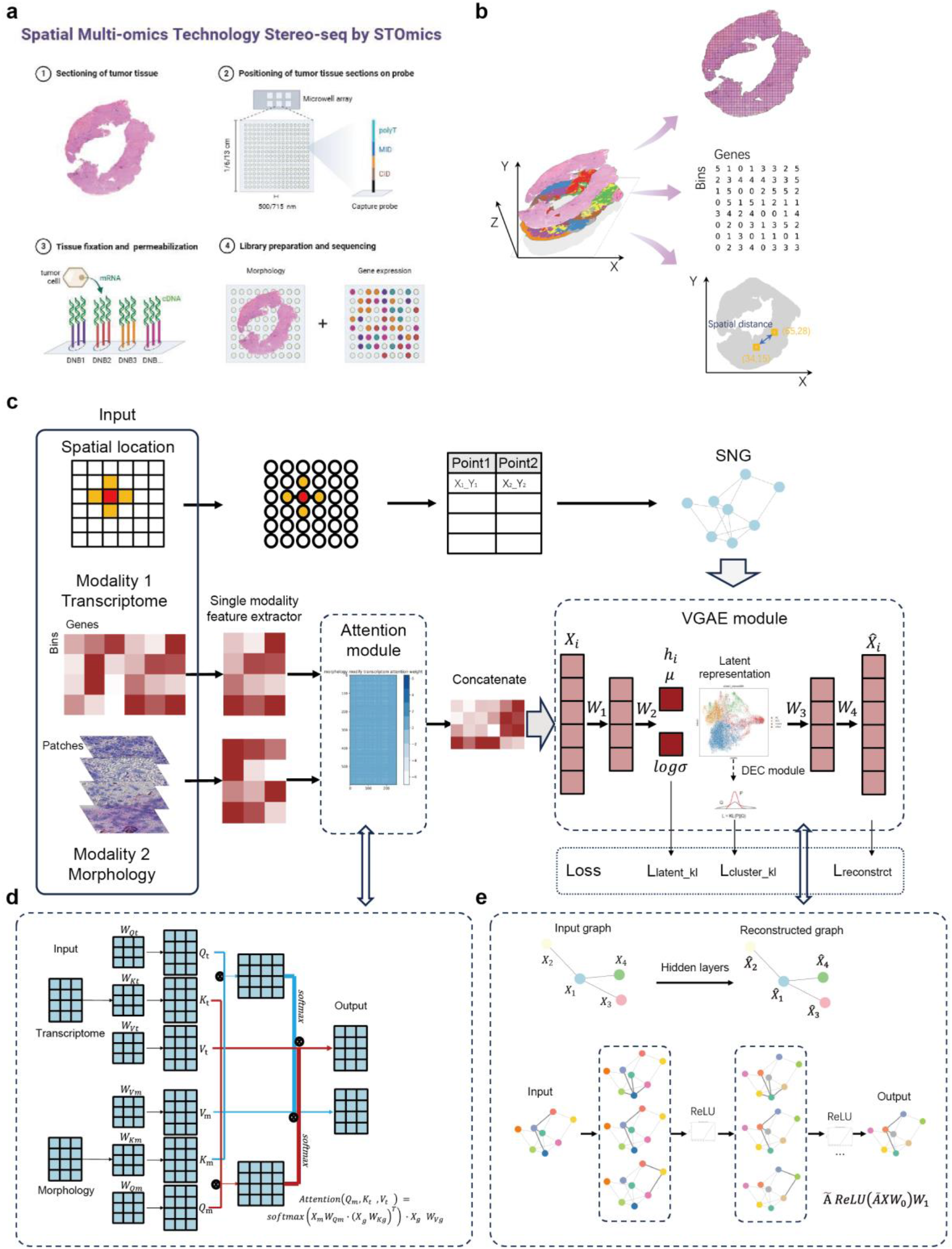
Schematic overview of StereoMM. **a**. Workflow for Stereo-seq experimental analysis. Created with BioRender.com. **b**. Data output formats from spatial transcriptomics. **c**. The overall framework of StereoMM. It requires three inputs: spatial coordinates, gene expression matrix, and image patches. Through the attention module and VGAE module, it generates low-dimensional a latent representation which can be used for downstream tasks. **d**. The cross-attention module in StereoMM captures relationships between different modalities by attending to relevant information from one modality based on another. In this module, each individual modality generates its own set of queries (Q), keys (K), and values (V). The Q from one modality is used to query the K and V from another modality. **e**. The VGAE module in StereoMM aggregates spatial information and each modality feature, and reduces the dimensionality of the original features through the encoder to obtain the final latent representation.

This framework utilizes a self-supervised Generative Neural Network (GNN) model (**Figure 2c**). It generates a feature representation that combines multiple modalities, which can be utilized for various downstream tasks to enhance the accuracy, such as spatial domain recognition. The learning process is guided by a combination of minimizing the self-supervised reconstruction loss and a regularization loss that forces the latent space representation. In an autoencoder, the reconstruction loss function promotes a high degree of similarity between the generated outputs 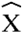 and the original input matrix (*X*), ensuring that the outputs closely mirror the inputs. In other words, it ensures that the latent features learned by the encoder preserve the maximum information from the original input, then the decoder can reconstruct the original input through these latent features. The intuition of the regularization loss, also known as the Kullback-Leibler (KL) divergence, encourages the model to learn a compact and smooth latent space representation.

Specifically, the training process is divided into the following four steps: (I) For the transcriptome and H&E image, a unimodal feature extractor is employed to extract *s*-dimensional unimodal features, generate two feature matrices (*X*_*t*_ ∈ *R*^*n×s*^ for transcriptome, and *X*_*m*_ ∈ *R*^*n×s*^ for morphology, where *n* represents the number of bins or spots). (II) These features are then fed into the attention module, where the information between modalities is integrated using the attention mechanism as in **Figure 2d**. This integration results in an *s*-dimensional output that enhances the interaction between modalities (*X*_*ta*_ ∈ *R*^*n×s*^ for transcriptome, and *X*_*ma*_ ∈ *R*^*n×s*^ for morphology). (III) The feature matrices from both modalities are concatenated (*X*=*X*_*ta*_ ⊕*X*_*ma*_) and used as input for the node features of the graph autoencoder. (IV) To incorporate spatial location information, a Spatial Neighbour Graph (SNG) is generated based on the physical distance. This SNG serves as the input for the adjacency matrix in the graph autoencoder.

The generative model for graph data utilizes the GNN to learn a distribution of node vector representations illustrated in **Figure 2e**. These representations are then sampled from the distribution, and the graph is reconstructed using the decoder. By extracting the latent representation from the Variational Graph AutoEncoder (VGAE), a high-quality, low-dimensional representation (*Z* ∈*R*^*n×d*^, Where *d* represents the feature dimension after dimensionality reduction) of the graph data is obtained. This feature representation Z can be effectively utilized for various downstream analyses, including clustering, trajectory analysis, and more.

### System parameter evaluation of StereoMM

We used a mouse brain tissues with intricate tissue structures as test sample for conducting a systematic evaluation of parameters. Firstly, we demonstrate that StereoMM outperforms individual modalities alone. We used anatomical reference annotations from the Allen Mouse Brain Atlas[15] as ground truth shown in **Figure 3a**. StereoMM accurately identified the hippocampal structure and differentiated mole and granul areas in the lobules shown by the rectangle in **Figure 3b**. In particular, StereoMM distinguished the subthalamic nucleus (domain 6), which is mainly composed of projection neurons and is a key part of movement regulation[16]. None of the single modality could independently identify this specific region (more details in **Supplementary Figure 1a**).

**Figure 3.**
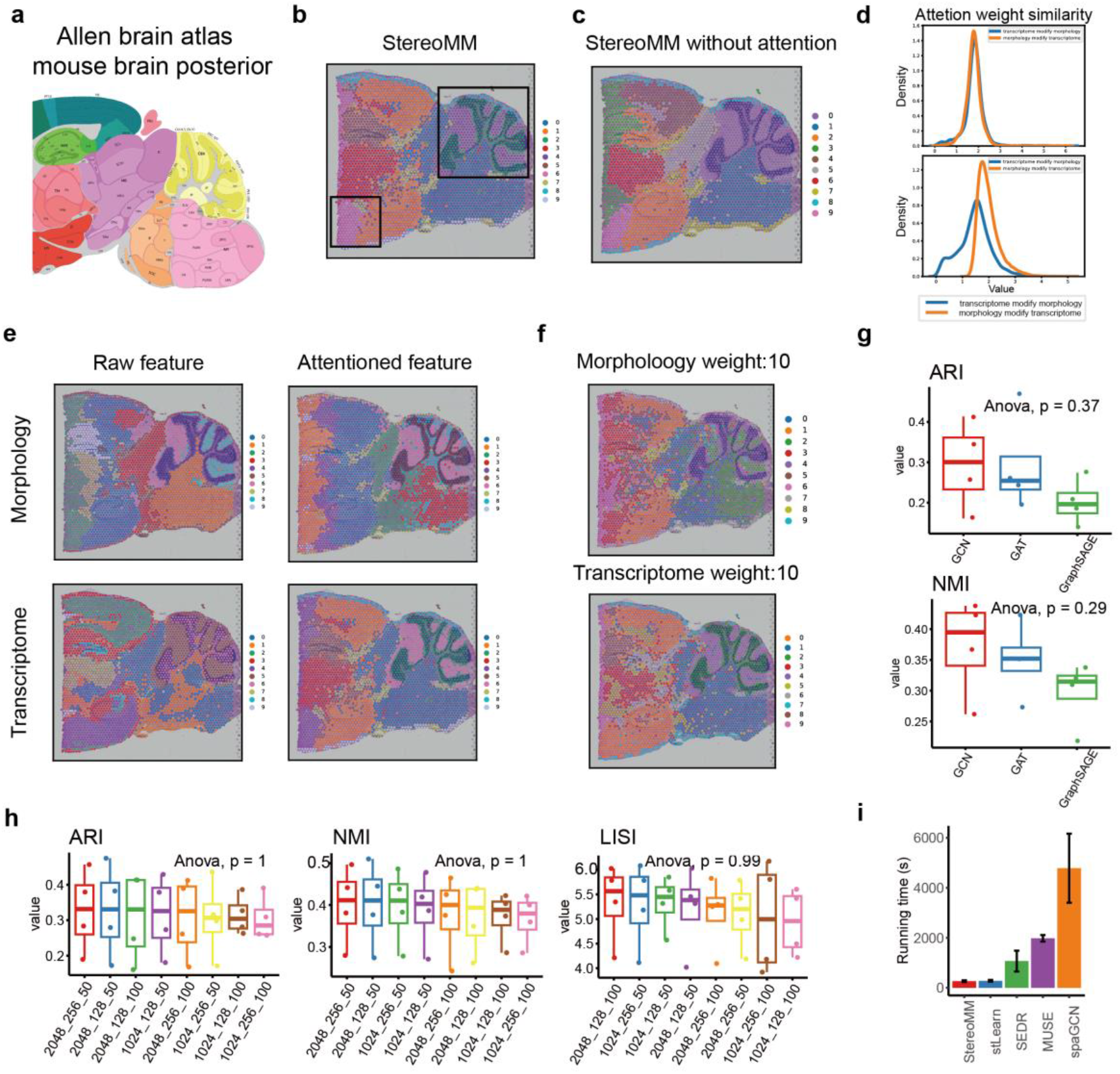
System parameter evaluation of StereoMM. **a**. The corresponding anatomical Allen Mouse Brain Atlas (https://atlas.brain-map.org/). **b**. Spatial domains identified by StereoMM. The black box denotes the cerebellar cortex and subthalamic nucleus. **c**. Spatial domains identified by StereoMM without attention module. **d**. The correlation between attention-enhanced features and final latent representations. On the top: the results on mouse brain slide, on the below: the results on slide3 of lung cancer. **e**. The features before and after the attention module are used to identify the spatial domain. **f**. The spatial domain recognized after manually setting the modality weight parameters. **g**. Boxplots of ARI and NMI values for three GNN types, each evaluated on 4 lung cancer slides. The center line, box lines, and whiskers of the boxplot represent the median, upper and lower quartiles, and 1.5× interquartile range, respectively. **h**. Boxplots of ARI, NMI and LISI values for different number of nodes per layer, each type evaluated on 4 lung cancer slides. **i**. Running time of 5 algorithms (StereoMM, stLearn, SEDR, MUSE, spaGCN) on all 4 lung cancer slices. The height of the histogram represents the average running time, and the whiskers represents the variance.

Therefore, we performed ablation experiments on the model to demonstrate the effectiveness of attention module. Without the interactive ability of attention, mole and granul areas in the lobules could no longer be distinguished, the identification of the hippocampal structure and subthalamic nucleus were also blurred, and more noise was introduced. Detailed comparisons are shown by the boxes in **Figure 3b** and **3c**. To clarify the role of the attention mechanism in enhancing explainability, we extract the weight matrix and compute its correlation with the final output (Z). This approach not only illuminates how our network assesses and assigns significance to individual modal features during fusion but also contributes to the model’s explainability by partially elucidating the decision-making process. In the mouse brain data, morphological similarity was on par with transcriptomic similarity, indicating that the model has fused the two aspects in a balanced manner. StereoMM also was tested on the lung cancer data of Stereo-seq, where the correlation between morphological features and the latent features is higher, suggesting that the model has assigned a higher weight to the morphological features in **Figure 3d**. In order to further illustrate the capabilities of the attention module, we extracted the features after the attention module for visualization in **Figure 3e**. After passing through the attention module, the two single-modal features become more similar and Mean Cosine Similarity increased from −26.76 to −13.91, indicating that the attention module enables mutual information exchange between the two modalities.

Meanwhile, we provided a hyperparameter to improve the guidance of prior knowledge. By setting custom weights for transcriptomic features, we maintain the flexibility of the model during the fusion process. As transcriptomic weight increased, the final output of the model became more similar to the transcriptome in **Figure 3f**. (ARI, from 0.17 to 0.29)

In the Stereo-seq lung cancer dataset, manual annotation in the pathology images, i.e. Whole Slides Imaging (WSI), served as the gold standard for quantification. We tested the impact of different hyperparameters on the accuracy of the results. StereoMM provided 3 types of convolutional neural networks for model selection, including Graph Convolutional Network (GCN), Graph Attention Networks (GAT) and Graph Sample and Aggregate (GraphSAGE). GCN achieved the optimal results in ARI and NMI as in **Figure 3g**. (More details in **Supplementary Figure 1b**). The user has a high degree of customization, with the ability to define the hidden layer of the network. For the model structure design, we assessed a total of 8 combinations of selecting 2048 or 1024 in the first layer, 256 or 128 in the second layer, and 50 or 100 in the third layer in **Figure 3h** and **Supplementary Figure 1c**. In general, StereoMM was robust under the choice of different number of nodes (ARI and NMI, Anova, p-value =1). However, an upward trend in ARI was observed with an increasing number of nodes, suggesting that the network’s enhanced fitting capability is due to the larger node count. 2048_256_50 achieved the highest average ARI score.

In demonstrating the model’s effectiveness and flexibility, we particularly highlighted its advantages in terms of running time. We conducted a comprehensive comparison of the time needed to execute various software tools, and our findings, as illustrated in **Figure 3i**, revealed that StereoMM required the shortest duration to complete its tasks. This efficiency underscores StereoMM’s superiority in processing speed, making it a highly practical choice for applications where time efficiency is critical.

Following detailed testing, the concept of the attention module showcased distinct benefits in terms of enhancing model performance and interpretability. Notably, it offered a clear method for adjusting weights for individual modalities within the attention module. Moreover, the StereoMM model’s architecture demonstrated resilience, efficiency, and accuracy under various parameter configurations.

### StereoMM improves performance of domain identification in Stereo-seq data of human lung adenosquamous carcinoma

To evaluate the accuracy of tissue identification and perform quantitative assessment of StereoMM, we conducted an analysis using lung adenosquamous carcinoma data generated from the Stereo-seq platform[17]. The data was meticulously annotated by pathologists into three distinct sections: lung adenocarcinoma (LUAD), lung squamous cell carcinoma (LUSC), and mixed areas in Figure 4a, which served as the gold standard. To reduce the computing burden, we divided the data into four slices (**Supplementary Figure 2a**). We also perform a benchmark analysis to compare the performance among each single modality, spaGCN, stLearn, MUSE and SEDR (**Figure 4b, Supplementary Figure 2b**).

**Figure 4.**
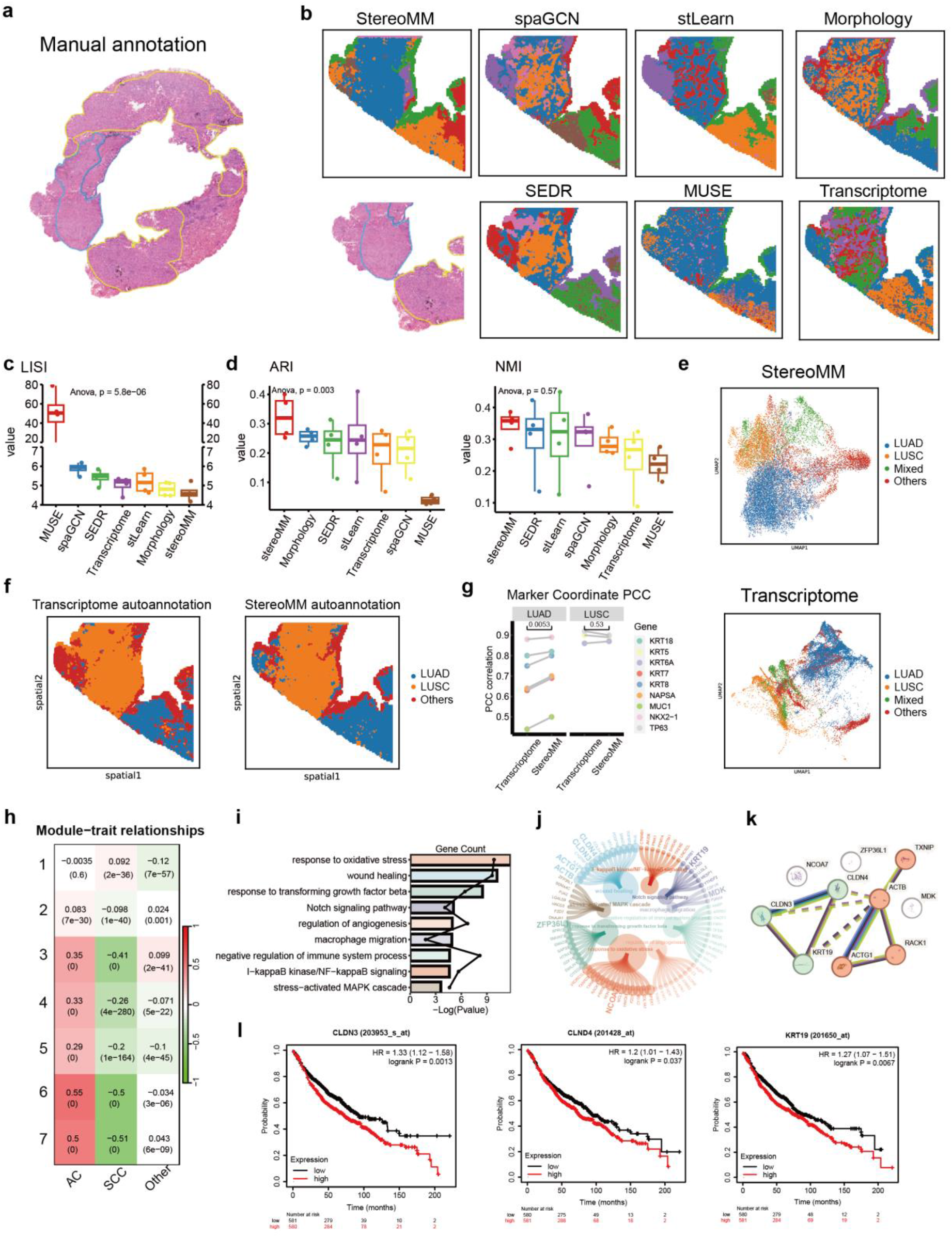
StereoMM improves recognition performance of human lung adenocarcinoma pathological regions. **a**. Manual annotation by pathologist. Area circled by red marker showed AC phenotype, blue displayed SCC phenotype. Green enclosed area presented mixed AC and SCC phenotypes. **b**. Manual annotation and the spatial domain identified by all algorithms on slice 4. **c**. Boxplot of LISI scores for seven methods in all 4 lung cancer slices. The center line, box lines, and whiskers of the boxplot represent the median, upper and lower quartiles, and 1.5×interquartile range, respectively. **d**. Boxplot of the cluster external evaluation index for seven methods in all 4 slices. **e**. UMAP visualizations of transcriptome and latent representation generated by StereoMM. **f**. Automated subtype annotation results from single transcriptome and StereoMM clustering. **g**. Spatial co-localization analysis of subtype annotations with corresponding marker genes. **h**. Heatmap of correlation between WGCNA gene modules and subtypes identified by StereoMM. **i**. GO functional enrichment results for Module 6. **j**. Circular visualization of genes within GO-enriched pathways. **k**. Protein-protein interaction network of hub genes. **l**. Correlation of genes within PPI clusters with prognosis.

We normalized the results of all methods to a consistent number of clusters (k=7). It has shown that StereoMM significantly enhances the accuracy of single-modal analysis. Single-modality features are noisy and exhibit a discontinuous distribution of clustering results. As expected, multimodal fusion significantly improved the issue of data noise (**Figure 4b**). To evaluate the noisy of the clustering results, we employed the local inverse Simpson’s index (LISI). A lower LISI score indicates better spatial separation. The LISI score of StereoMM is 4.64±0.44, which is lower than that of single transcriptomic features (4.81±0.37) or single morphological features (5.02±0.44). Which demonstrated that StereoMM achieves superior spatial separation compared to methods that rely solely on transcriptomic or morphological features (**Figure 4c**).

In addition, compared with previous spatial clustering methods that combined histology or spatial, StereoMM exhibits significant improvement in spatial recognition ability. We quantitatively assessed its capabilities using several indicators, including evaluation metrics with the gold standard of fundamental organizational facts: Adjusted Rand Index (ARI) and Normalized Mutual Information (NMI) (**Figure 4d**). Furthermore, internal evaluation metrics of clustering are calculated. These metrics provide insights into the quality and performance of clustering results by measuring the separation and compactness of clusters. The commonly used internal evaluation metrics including: Calinski-Harabasz Index (CH), Davies-Bouldin Index (DB), Silhouette Coefficient (SC) (**Supplementary Figure 3a**). Except for the DB score, higher scores in all the mentioned metrics indicate better performance. We also calculated LISI scores for all methods (**Figure 4c**). Except for CH, StereoMM achieved the best performance, obtaining the highest ARI (0.32±0.07) and NMI (0.34±0.05), demonstrating its exceptional performance in accurately identifying different tissue types. We visualized the embeddings of StereoMM using UMAP graphs. Comparing the distribution of the original transcriptome, StereoMM clearly separated the three manually annotated categories, while the original transcriptome showed a mixed and disordered state (**Figure 4e, Supplementary Figure 3b**).

To validate the enhanced accuracy of StereoMM in identifying clinical regions compared to single transcriptomics (**Figure 4f**), we selected commonly used clinical diagnostic markers for LUAD (NKX2-1, KRT7, NAPSA, MUC1, KRT8, and KRT18) and LUSC (KRT5, KRT6A, TP63)[18] (**Supplementary Figure 4a**), and then quantified the spatial co-localization with each molecule using the Kernel Density Estimation (KDE) and Pearson Correlation Coefficient(PCC)[19] (**Supplementary Figure 4b-c**). We found that the accuracy of StereoMM is greater than single transcriptomics, as evidenced by the higher correlation between the automated annotation results of StereoMM and the molecular expression of LUAD(P=0.0045<0.05). However, for LUSC, the comparison of identification accuracy between both methods was not statistically significant (P=0.67>0.05) (**Figure 4g**). Subsequently, we used weighted gene co-expression network analysis (WGCNA) to cluster gene expression into seven modules and calculated the spatial correlation of these modules with StereoMM annotated regions (**Figure 4h, Supplementary Figure 5a-c**). Only Module 1 was linked to LUSC, but Modules 2–7 had a stronger association with LUAD. Module 3 exhibits the strongest association with other regions aside from this. We concluded that Module 5, which showed the highest correlation with LUAD, was functionally biased toward immunosuppressive and tumor growth after performing gene enrichment analysis (GO&KEGG) (**Figure 4i**). The pathways associated with macrophage migration (CSF1R[20]), NF-κB signalling (CTNNB1[21]) and TGF-β signalling (SMAD5[22]) were found to be overexpressed (**Figure 4j**). We referred to the genes within module 5 that interacted more with other genes as eigengene genes and matched them with corresponding pathways. After conducting a protein-protein interaction (PPI) network analysis of these hub genes, we discovered that the interaction between CLDN3 (claudin 3), CLDN4 (claudin 4), and KRT19 (cytokeratin 19) was the most significant exclude irrelevant genes (**Figure 4k**), suggesting that these might be important genes affecting the function of Module 5. CLDN3 and CLDN4 are tight junction molecules correlated with ovarian cancer cell infiltration and wound healing[23], while KRT19 is a member of the keratin family and related to Notch pathway[24]. All three are overexpressed in lung adenocarcinoma and are associated with epithelial-mesenchymal transition (EMT) and tumor metastasis. Therefore, we calculate the association of these genes with the prognosis of LUAD patients via the Cancer Genome Atlas (TCGA) database. The results exhibited higher expression of CLDN3, CLDN4, and KRT19 was associated with poor prognosis in LUAD patients (**Figure 4l**), indicating that these molecules may promote tumor development and be the potential biomarkers[25-27].

**Figure 5:**
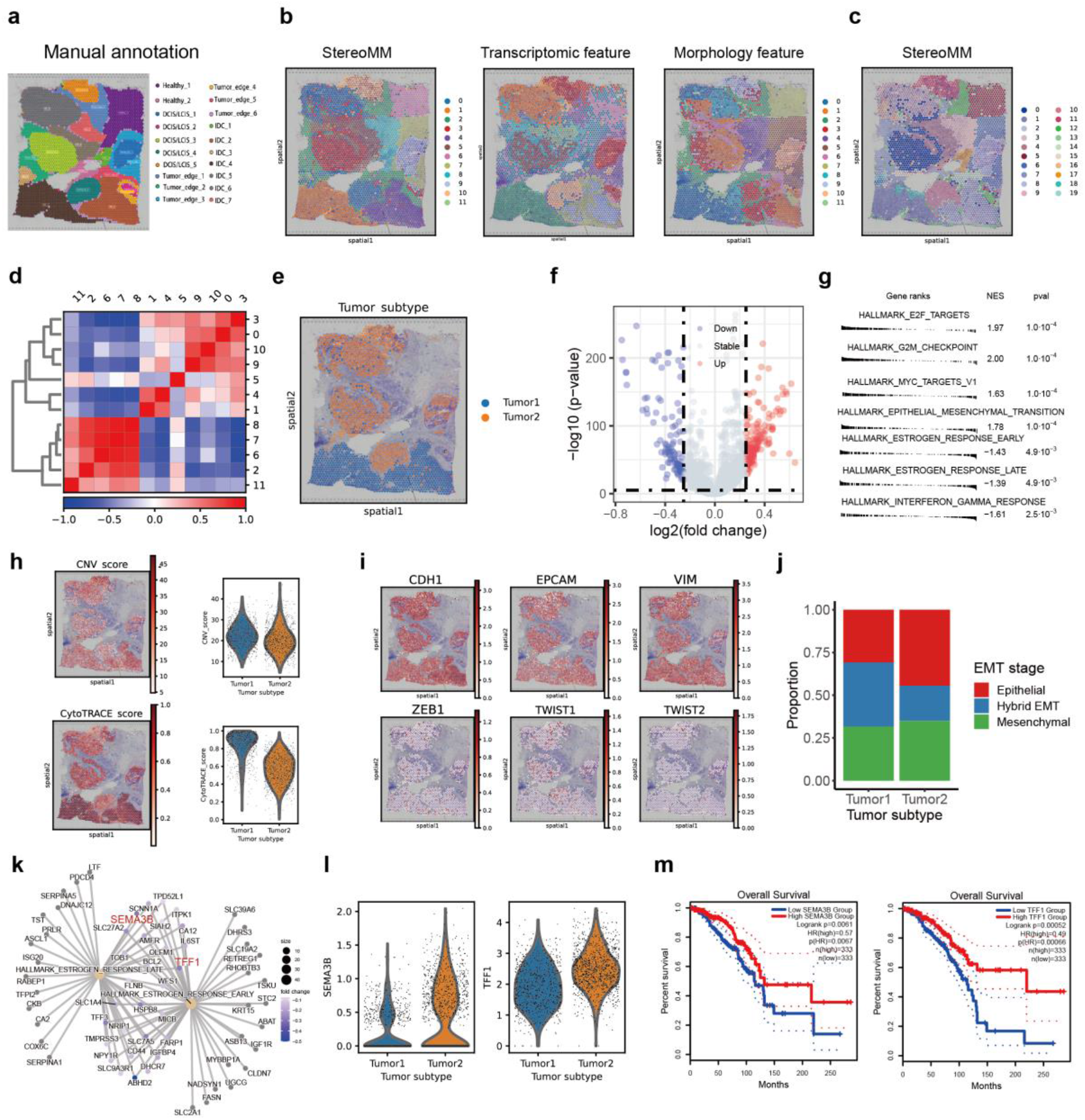
StereoMM dissects breast cancer heterogeneity. **a**. Manual pathological annotation based on hematoxylin and eosin staining of human breast cancer data. IDC, invasive ductal carcinoma; DCIS, ductal carcinoma *in situ*; LCIS, lobular carcinoma *in situ*; tumor edge; healthy region. **b**. Spatial domains identified by StereoMM (left) and each single modality (middle: single transcriptome; right: single morphology). **c**. Spatial domains identified by StereoMM with 20 clusters. **d**. Heatmap of pearson correlation coefficient between domains (domains= 12). **e**. Volcano plot visualization of DEGs between tumor1 and tumor2. **f**. GSEA showed related pathways enriched in different tumor subtypes (tumor1 and tumor2). **g**. CNV scores and differentiation calculated by CytoTRACE for different tumor subtypes. On the left: visualization of spatial location of CNV scores. On top right: CytoTRACE scores for different tumor subtypes. On bottom right: CNV scores for different tumor subtypes. **h**. Spatial location of the expression of EMT-related marker genes. **i**. Proportion of EMT status in different tumor subtypes. **j**. Potential gene regulatory network of estrogen response pathway (early and late). **k**. Expression levels of genes shared by estrogen response pathways (SEMA3B and TFF1) in different tumor subtypes. **l**. Survival curves of SEMA3B and TFF1 genes in TCGA breast cancer database.

In summary, the architecture based on attention and graph neural networks used by our structure helped capture and combine information that could not be obtained from either mode alone. A fair comparison of results showed that the recognition ability in the spatial domain of StereoMM was significantly better than that of a single modality or any competing software, whether based on gold standard indicators or other indicators. Simultaneously, StereoMM can assist in identifying significant genes and putative targets related to the initiation and progression of tumors.

### StereoMM dissects breast cancer heterogeneity and identifies potential prognostic factors

To assess the capability and compatibility of StereoMM, we applied StereoMM to an open-access dataset generated from the fresh frozen invasive ductal carcinoma breast tissue using the 10x Visium spatial platform. For this dataset,StereoMM not only revealed the clear clustering structure which was consistent with the manual annotation, but also specifically identified tumor boundary area as a separate domain **(Figure 5a-b)**. Next, we increased the number of clusters to validate the robustness of StereoMM, and successfully distinguishing separate tumor boundary regions, DCIS/LCIS regions, as well as the smallest IDC region **(Figure 5c)**.

To further investigate the intricate tumor microenvironment and explore the biological characteristics within different spatial compartments, we performed a correlation analysis between the domains identified by StereoMM (domains=12), and discovered the tumor area was divided into two parts which was not completely consistent of histological phenotype (**Figure 5d-e**). We first focused on comparing intratumoral transcriptional differences between tumor1 (including domain1 and 4) and tumor2 (including domain 0,3,9 and 10) by performing differential expression analysis followed by gene set enrichment analysis (GSEA). We detected significant DEGs (|log fold change| ≥0.25; p-value < 0.05) between tumor 1 and 2 (**Figure 5f**). In tumor1, (Figure 5f) ‘E2F_TARGETS’, ‘G2M_CHECKPOINT’ and ‘EPITHELIAL_MESENCHYMAL_TRANSITION’ pathway were upregulated, while ‘INTERFERON_GAMMA_RESPONSE’, ‘ESTROGEN_RESPONSE_LATE’ and ‘ESTROGEN_RESPONSE_EARLY’ were downregulated (Figure 5g). These pathways can interact with each other and are associated with the prognosis and treatment response of breast cancer[28, 29].

To specifically assess the heterogeneity between tumor1 and tumor2, we next performed copy number variation (CNV) analysis and differentiation analysis using inferCNV and CytoTRACE respectively (**Figure 5h**), and described the different EMT tumor states based on the expression of E-cadherin (E-cad) and vimentin (VIM). As expected, tumor1 displayed a distinctively higher inferCNV score(t-test, p-value = 6.44e-12)and CytoTRACE score (t-test, p-value = 5.69e-236), indicating the heterogeneity of tumor proliferation and malignancy. Then we investigated the expression of EMT markers (**Figure 5i**), including epithelial molecules (E-Cadherin and EPCAM), mesenchymal markers (VIM) and transcription factors associated with EMT (ZEB1, TWIST1 and TWIST2). Next, we annotated tumor epithelial cell by deconvolution and cell2location (**Supplementary Figure 6a**), and then defined distinct EMT cell state ranging from epithelial (E-cad+ VIM-), hybrid EMT (E-cad+ VIM+) and mesenchymal (E-cad-VIM+)[30, 31] (**Figure 5j**). We observed tumor1 increased the proportion of the hybrid EMT and decreased the proportion of epithelial, indicating the possibility of infiltration and metastasis. On the other hand, GSEA results displayed different estrogen response across regions (**Figure 5k**), which is relevant to the published clinical information of the sample (ER+PR-HER2+). Meanwhile, we observed upregulation of SEMA3B and TFF1 in tumor2 (**Figure 5l**), which tend to exhibit tumor suppressor function and are reported as potential biomarkers in breast cancer (BC) before. We also validated the function of SEMA3B and TFF1 using survival data from TCGA cohort of 333 HER2+ BC patients (**Figure 5m**), suggesting the prognosis value of SEMA3B and TFF1.

Another interesting finding is that in the correlation analysis with 12 cluster, domain11 initially labelled as IDC was clustered with healthy tissue. The DEG and GSEA results indicate upregulated oncogenic pathways, immune-related pathways, and B-cell markers in this domain (**Supplementary Figure 6b-c**), suggesting the potential presence of tertiary lymphoid structures. This is consistent with previous studies[32] (**Supplementary Figure 6d**).

In summary, analysis of StereoMM clusters revealed regional and biological differences reflecting tumor progression and raised the hypothesis that heterogeneity of proliferation and differentiation states result the distinct capability of metastasis and resistance to therapy across histologic subtypes.

## Conclusion and Discussion

The amalgamation of histopathology with high-throughput sequencing to inform oncologic treatment strategies is in its infancy. Spatial omics has emerged as a powerful tool in precision medicine, outperforming established metrics such as tumor mutational burden in predicting responses to PD1/PD-L1 therapies in a pivotal clinical trial[1]. Nevertheless, the utility of spatial transcriptomic data is constrained by limitations such as low total transcriptions per cell, significant data noise, and a high frequency of zero values, necessitating the integration of additional modal data for a comprehensive analysis[2-4]. Thus, the innovation of effective modal fusion methodologies is imperative.

Several algorithms have been designed to integrate information from MM of the ST data. stLearn is a widely used spatial transcriptomics analysis tool. However, it does not perform appropriate weighting when normalizing using histological images with spatial location. spaGCN utilizes graph convolutional neural networks (GCN) to model spatial relationships[33]. While, it has limited capabilities in feature extraction because it simply utilizes the pixel values of the three channels of the image and ignores the high-level features of morphology. Software such as MUSE[34] and SEDR employ architectures underpinned by autoencoders to learn a low-dimensional representation of multimodal data, but such integration relies entirely on neural networks and lacks interpretability. While these methods have yielded numerous intriguing findings, they may be limited in their flexibility, generality, and the interpretability of model decisions. These limitations can hinder their application in real-world projects.

Our study introduces StereoMM, a deep learning approach that integrates multimodal data—including high-content H&E images, spatial information, and gene expression—to comprehensively identify tumor subpopulations, significantly advancing beyond conventional methods by considering both histological and cellular interactions within tissue samples. StereoMM employs an attention mechanism for deep interaction between modalities, followed by the aggregation of multimodal features from adjacent tissues using a graph convolutional network. This methodology affords StereoMM with exceptional adaptability and computational efficiency. The utility of the attention module in mediating information exchange has been substantiated through ablation studies and similarity assessments. By adjusting various parameters, we have demonstrated the robustness of our model, which does not preclude users from fine-tuning based on their understanding of the data. For instance, tissues with lower inter-regional similarity may benefit from a smaller k-nearest neighbours parameter or fewer graph convolutional layers. Such customization can yield results with greater biological relevance across diverse datasets.

StereoMM has been validated on tumor datasets from Stereo-seq and 10X Visium, exhibiting superior performance in spatial contour identification. Comparative analyses with manual annotations have revealed spatial domains that more accurately reflect the ground truths, and congruence with cell subtype marker genes has indicated subpopulation compositions that correlate with biological functions. The intricate spatial architecture of tumor tissues necessitates a detailed analysis of the spatial microenvironment, which is crucial for comprehending tumor biology, unravelling mechanisms of oncogenesis, and identifying therapeutic targets. The refined subpopulations discerned through StereoMM, in conjunction with multimodal data, appear to capture significant biological variations, including genes implicated in tumor progression and intratumoral heterogeneity.

At present, StereoMM has been applied to spatial transcriptomic analyses using binning or meshing methods. While the modeling framework of StereoMM is theoretically applicable to other spatial transcriptomics platforms, the rapid evolution of ST technology presents new measurement techniques[5, 6], expanded data volumes, and progress in additional modalities[7]. Consequently, the development of novel methods to exploit the expanding spatial transcriptomic data represents a considerable challenge. The scalability of the model can be enhanced through strategies such as subgraph sampling and parallel training. Moreover, the incorporation of non-aligned modal data from beyond spatial transcriptomics could bolster our capacity to analyse and interpret tissue heterogeneity. Future investigations will explore these potential enhancements to further refine the functionality of StereoMM.

In summary, StereoMM is an innovative and promising approach utilizing attention mechanisms and graph autoencoders for the analysis of spatial transcriptomic data. It facilitates modality fusion through self-supervised learning in the absence of annotations. Poised to capitalize on forthcoming advancements in measurement technologies, StereoMM holds the potential to significantly improve precision oncology practices in the context of therapeutic decision-making.

## Acknowledgement

The author would like to acknowledge Mr. Zidong Su (BGI Research Institute, Shenzhen) for his assistance in the design of the theoretical framework. This study is supported by Science and Technology Innovation Key R&D Program of Chongqing (CSTB2023TIAD-STX0002).

## Ethics Approval and consent of participate

This study does not require ethics approval or informed consent from participants.

## Competing Interests

The authors declare no competing interests.

## Consent for Publication

The authors declare that the research was conducted in the absence of any commercial or financial relationships that could be construed as a potential conflict of interest.

## Code Availability

Code for data analysis is available at https://github.com/STOmics/StereoMMv1.

## Disclosure/Competing Interest

The authors declare no potential competing interests.

## Author Contributions

Conceptualization: Bingying Luo, Jiajun Zhang, Xun Xu, Ao Chen and Fei Teng.

Project administration and supervision: Jiajun Zhang, Xun Xu, Ao Chen, Sha Liao, Xi Feng and GuiBo Li.

Soft development and implementation: Bingying Luo and Fei Teng.

Data collection, processing, and application: Bingying Luo, Fei Teng, Jiajun Zhang, WeiXuan Chen, Mei Li, Xuanzhu Liu, Huaqiang Huang, Yu Feng, Xing Liu, Min Jian, Xue Zhang.

Method comparisons: Bingying Luo, Xuanzhu Liu.

Manuscript writing: Bingying Luo, Guo Tang, Jiajun Zhang, Fei Teng.

Figure generation: Bingying Luo and Fei Teng.

Manuscript review: Jiajun zhang, XunXu, Feng Xi, Guibo Li, Qu Chi, Xin Liu.

Project coordination: Jiajun Zhang, Fei Teng, Sha Liao, and Ao Chen.

Biological interpretation: Bingying Luo, Guo Tang, Jiajun Zhang, and Fei Teng.

Manuscript review: Jiajun Zhang, Xun Xu, Ao Chen, Sha Liao.

## Notes

### Competing Interest Statement

The authors have declared no competing interest.

