## Supplemental Figure, Materials And Methods for "StereoMM: A Graph Fusion Model for Integrating Spatial Transcriptomic Data and Pathological Images"

#### Image registration scheme

For Stereo-seq data, the registration operation in StereoMap is used to align morphological images and spatial transcriptomics data. We choose the manual registration approach. Firstly, click on the manual registration tool button to open the manual registration mode. Then, in the registration layer, select the image that needs to be registered. Secondly, feature points can be identified, and the image can be translated, rotated, and normalized around these feature points. Finally, the registration is completed by aligning the feature points and applying affine transformations, stretching, etc., to the image. After the registration, the morphological image and spatial transcriptome data are corresponded in spatial coordinates and can be aligned for feature fusion. The resolution size of the registered image is the same as the bin1 data size of Stereo-seq.

#### Data description and preprocessing

We used three datasets to test the capability of StereoMM. These datasets include a lung cancer dataset from the Stereo-seq platform, a breast cancer dataset from the 10X platform, and two mouse brain datasets from the 10X platform. The H&E images of all datasets were registered with the transcriptomic coordinates.

For the lung cancer dataset from the Stereo-seq platform, we divided it into four slices. Each slice was aggregated in a non-overlapping manner with a resolution of 100*100 DNB (bin100) to generate the initial expression matrix. The scanpy package was used to construct anndata for downstream analysis, and cells with a total count less than 300 were filtered out. For the 10X platform datasets, the raw data was downloaded from the database and then directly read into anndata using the read_visium function in scanpy. Spots with in_tissue=0 were filtered out. After obtaining the anndata for all the datasets, the same preprocessing steps were applied to the expression matrices: data normalization were performed using scanpy with normalize_total, log1p and scale functions. Unsupervised clustering is performed using the tl module function within the scanpy API.

### The StereoMM model

StereoMM is a self-supervised learning model that requires three profiles: gene expression matrix, morphological image and spatial location information. The ability of StereoMM to combine multimodal data is built upon the following two principles: (1) fully integrating features between modalities, (2) faithful preservation of effective information from original modalities during dimensionality reduction. These two principles are respectively implemented by the attention module and the graph autoencoder module. Additionally, in order to enable the model to have the ability of joint analysis of multiple slices and optimized clustering, we also provide the deep embedded clustering (DEC) module. The specific training process for StereoMM is as follows:

(1) Generate tiled images based on the center points of each bin or spot. The high-level visual features of each patch ( $M = \{M_1, M_2, \cdots, M_n\}$) are learned through an imaging single-modal feature extractor, convolutional neural network (CNN) and vision transformer (VIT) methods are two optional methods. The output before the last fully connected layer is extracted as the image feature of an image patch. Each patch can obtain an *s*-dimensional image feature vector (for ResNet50, $s = 2048$ , $x_{mi} = \left[vec(M_{i,1}), vec(M_{i,2}), \cdots, vec(M_{i,s})\right]$). For the entire slide data, we can obtain a morphological feature matrix ($X_m \in R^{n \times s}$). Where *n* is the number of bins or spots.

(2) For the gene expression matrix, we obtain the features of each bin or spot ($T = \{T_1, T_2, \cdots, T_n\}$) through a single-modal feature extractor (principal component analysis, highly variable genes or genes with highest variance). We maintain the dimensionality of transcriptomic features consistent with the dimensionality of morphological features ($x_{ti} = [vec(T_{i,1}), vec(T_{i,2}), \cdots, vec(T_{i,s})]$). For the entire slide data, we can obtain a transcriptomic feature matrix ($X_t \in R^{n\times s}$).

(3) Once the transcriptome and morphological features are obtained, a cross-attention module is used to fuse each other features from two different modal, ensuring that the output feature dimension remains consistent with the initial dimension. The feature dimension after attention ($s = 2048$) remains the same as before, but the feature vector updates.

(4) Updated transcriptomic and morphological features are concatenated as node representations, which together with the adjacency matrix of SNG constitute the final graphical representation: $G = (V, E)$. Where $v_i \in V$ represents the $i$ th bin or spot, and $i = 1, 2, \cdots, n$ represents all $N$ bins or spots. $e_{ij} \in E$ represents the connection between $v_i$ and $v_j$. The graph is reconstructed through the variational graph autoencoder, and the intermediate latent space is extracted as a new fusion feature representation ($Z \in R^{n\times d}$). Where $d$ represents the feature dimension after dimensionality reduction. The new fused feature representation $Z$ can be used for downstream analysis tasks, here we focus on clustering.

StereoMM offers an optional fifth step: operation DEC after extracting the latent space. DEC leverages deep clustering theory to provide the model with optimized features for clustering problems and the ability to perform joint analysis across multiple slices.

**Construction of spatial neighbour graph (SNG)**

To aggregate the information of neighbouring points for a specified point, StereoMM generates an undirected neighbour graph called spatial neighbour graph (SNG) based on spatial coordinates.

There are two methods to construct the SNG using the nearest neighbour algorithm: (1) the number of the nearest neighbours, (2) the radius threshold. In method 1, the bins or spots (hereinafter referred to as points) are sorted based on the Euclidean distance between the specified point and other points, and the k nearest points to the specified point are selected. In method 2, all points within a certain Euclidean distance from the specified point are chosen as neighbours. For all data in this article, a radius threshold is used to construct the SNG. Specifically, for Stereo-seq data with bin100, a radius threshold of 100 is chosen. For 10X Visum data, the distance between each spot is calculated based on the resolution of the full image, and this distance is used as the threshold.

Through the above calculation, assuming $A$ is the adjacency matrix of SNG, then $A_{ij} = A_{ji}$ are assigned a value of 1 when there is a connection between vertex $i$ and vertex $j$, otherwise it is assigned a value of 0.

By using this approach, we can construct a neighbour graph that includes the nearest neighbouring cells of a specified point, allowing for the aggregation and analysis of neighbouring cell information. This method helps us better understand and interpret the cell-cell relationships and interactions in spatial transcriptomics data.

### Attention mechanism

The attention mechanism module is the main component of the StereoMM model. The attention mechanism has the capability to automatically learn the information in the query and selectively focus on important information, thereby enhancing the performance of information interaction. The attention mechanism utilizes three main components: queries (Q), keys (K), and values (V).

For the morphology-dependent transcriptomic attention mechanism, we perform the following operations:

StereoMM utilizes three fully connected layers to map transcriptomic features to key tensors and value tensors, as well as map morphology features to a query matrix.

$$Q_m = X_m W_{Qm}$$

$$K_t = X_t W_{Kt}$$

$$V_t = X_t W_{Vt}$$

Where $W_{Qm}, W_{Kt}, W_{Vt} \in R^{n \times d_k}$. Here, $d_k$ denotes the inner dimension of that is specific to each attention layer.

We compute morphology-dependent transcriptomic attention weights by the following steps:

Step 1: Calculate the inner product to determine the correlation between $Q_m$ from morphology and $K_t$ from transcriptomics.

$$simi\left(K_t, Q_m\right) = Q_m \bullet K_t^T$$

Step 2: The similarity is normalized by softmax to obtain the correlation weight between morphology and transcriptome.

$$\alpha_{K_t, Q_m} = softmax\ (simi(K_t, Q_m)) = \frac{\exp\left(simi\left(K_t, Q_m\right)\right)}{\sum_{i=1}^{n} simi\left(K_t, Q_m\right)}$$

Step 3: Calculate the final attention weight through value and similarity score

$$Attention\ (Q_m, K_t, V_t) = \alpha_{K_t, Q_m} \bullet V_t$$

Finally, we can define the attention layer as:

$$Y = softmax(X_m W_{Qm} \bullet (X_g W_{Kg})^T) \bullet X_g W_{Vg}$$

The process of setting the transcriptome as the query involves similar operations. The key difference is that the query is derived from the transcriptome feature matrix, while the key and value are derived from the morphological feature matrix.

#### Graph variational autoencoder

Graph variational autoencoder (VGAE) is used to obtain an embedded representation of multimodal data that incorporates SNG. The autoencoder structure can be guided by self-supervised methods. A standard autoencoder consists of an encoder and a decoder.

Replacing the linear layers of the standard autoencoder with graph neural networks can incorporate spatial information. And variational modifications to standard encoder-decoder architectures can be transformed into probabilistic encoders, improving the performance of spatial embeddings.

Given the cell adjacency matrix $A$ and the node feature matrix $X$ ($X \in R^{n \times 2s}$) obtained by concatenating morphological features and transcriptome features. VGAE learns the latent representation $Z$ ($Z \in R^{n \times d}$)of the cell graph. Then, $Z$ is used to reconstruct the node feature matrix or adjacency matrix in the encoder.

The generative model (encoder) is divided into the following two parts:

Step1: Get the mean ($\mu$) and the logarithm of the variance ($log\ \sigma$). The inference part of VGAE is parameterized by graph neural networks (GNNs). StereoMM provide three options for the GNN architecture, including GCN, GAT, and GraphSAGE. By default, StereoMM use GCN.

$$\mu = GCN_{\mu}(X, A) = \tilde{A}\ ReLU(\tilde{A}XW_{\mu 0})\ W_{\mu 1}$$

$$log\sigma = GCN_{log\sigma}(X, A) = \tilde{A}\ ReLU(\tilde{A}XW_{log\sigma 0})\ W_{log\sigma 1}$$

$$\tilde{A} = D^{-\frac{1}{2}} A D^{-\frac{1}{2}}$$

Where, $W_{\mu i}$ and $W_{log\sigma i}$ respectively represent the weight matrix of $i$ th layer used to infer $\mu$ and $log\ \sigma$, and $\tilde{A}$represents the symmetrically normalized adjacency matrix. By default, the first layer and the second layer are set to 2048 and 512, respectively.

Step2: According to the $\mu$ and the $log\ \sigma$, obtain the hidden representation $Z$.

$$q(Z \mid X, A) = \prod_{i=1}^{N} q(z_i \mid X, A)$$

$$q(z_i \mid X, A) = N(z_i \mid \mu_i, diag(\sigma_i^2))$$

The variational approximation (decoder) reconstructs node features using hidden representation of the same GNN type as the encoder. Taking GCN as an example, the decoder includes i+1 layers.

$$\widehat{X} = \tilde{A}\Big(ReLU\Big(\tilde{A}(ReLU\Big(\tilde{A}ZW_{d0}\Big)W_{d1}\Big)W_{d2}\Big)$$

Where $W_{di}$ represent the weight matrix of $i$ th layer in decoder.

For different task requirements, although it is not the default setting, we also provide a decoder that reconstructs the graph structure.

$$p(A \mid Z) = \prod_{i=1}^{N} \prod_{j=1}^{N} p(A_{ij} \mid z_i, z_j)$$

$$p(A_{ij} = 1 \mid z_i, z_j) = \sigma(z_i^T z_j)$$

where $A_{ij}$ are the elements of $A$ and $\sigma(\cdot)$ is the logistic sigmoid function.

**Loss function**

StereoMM uses VGAE to reconstruct graphs and perform self-supervised training. Generally, the model's loss consists of two parts: (1) Reconstruction loss: the error when reconstructing node features, this component measures the ability of VGAE to reconstruct nodes. (2) Variational Kullback-Leibler (kl) divergence loss: the error between the variational probability distribution and the prior distribution. VGAE uses variational inference to learn the latent variable distribution of the nodes and maximize the lower bound estimation of the node representations. The loss is typically computed using the kl divergence between the learned distribution and the prior distribution of the latent variables. Ultimately forces two distributions to be similar and provides regularity.

(1) Reconstruction loss:

$$L_{recon} = MSE(X, \widehat{X}) = \frac{1}{n} \sum_{i=1}^{n} \left( X_i - \widehat{X}_i \right)^2$$

Where $\widehat{X}$ represents the node feature matrix reconstructed by the decoder of VGAE.

(2) Variational kl divergence loss:

$$L_{vkl} = KL(q\ (Z|X, A))\, ||p(Z) = \frac{1}{2} \sum_{i=1}^{n} \left( 1 + log\sigma_i^2 - \mu_i^2 - \sigma_i^2 \right)$$

the total loss function of the SGATE model is summarized as:

$$L_{total} = L_{recon} + \beta L_{vkl}$$

Where $\beta$ is a parameter that controls the weight of the two losses, the default is $\frac{1}{n}$. And n is the number of bins or spots.

**Optimization by DEC**

We offer an optional deep embedding cluster (DEC) step. The DEC algorithm iteratively optimizes the autoencoder and cluster centers to continuously learn the data representation and clustering results, thereby achieving better clustering performance. In addition to providing a better representation for clustering, DEC also has the ability to eliminate batch effects.

**StereoMM parameter settings**

For the H&E images, we segmented the entire chip region based on each bin or spot (hereinafter referred to as point) centers of transcriptome. This will produce a consistent number of image patches with the number of points in the transcriptome. For the bin100 data from Stereo-seq, we used patches of size 128*128 pixels (64*64 μm) corresponding to the centers of the corresponding transcriptomes. For the spot data from 10X, patches of size 200*200 pixels were extracted by the same way. Morphological features were extracted from each patch using the default parameters of a pre-trained ResNet50 model.

When performing the fusion strategy with StereoMM, consistent parameters were used for all the datasets: GNN with GCN architecture and a hidden layer structure of 2048-512-100.

**Spatial domain detection**

After applying StereoMM to analyse ST data, we learn accurate low-dimensional latent representations ($Z$) that represent the multimodal fused features of each bins or spots. StereoMM uses Leiden algorithm to cluster the latent representations of the bins or spots. In the Leiden algorithm, the graph is typically constructed by connecting each node to its k nearest neighbours. The selection of neighbours is crucial as it defines the local neighbourhood structure used for clustering. For fair comparison, we take the same default 20 nearest neighbours for all data to build the graph. We adjust the value of resolution to determine the final number of clusters. These clustered groups are defined as spatial domains.

### Spatial domain evaluation

We employ multiple different metrics to evaluate the clustering results of spatial domains. These metrics include both internal and external indicators of clustering quality, as well as measures that assess spatial diversity.

### Clustering external indicators

For data where true sub-group labels are available, we use external clustering metrics for evaluation. Adjusted Rand Index (ARI) and Normalized Mutual Information (NMI) are widely used external clustering metrics to evaluate the consistency between clustering results and external labels or reference information.

$$AdjustedIndex = \frac{Index - ExpectedIndex}{MaxIndex - ExpectedIndex}$$

$$ARI = \frac{\sum_{ij}\binom{n_{ij}}{2} - \left[\sum_i \binom{a_i}{2} \sum_i \binom{b_j}{2}\right] \Big/ \binom{n}{2}}{\frac{1}{2}\left[\sum_i \binom{a_i}{2} + \sum_j \binom{b_j}{2}\right] - \left[\sum_i \binom{a_i}{2} \sum_j \binom{b_j}{2}\right] \Big/ \binom{n}{2}}$$

ARI measures the similarity between clustering results and external labels. It takes into account the impact of randomness on the Rand Index and adjusts it to have a range between -1 and 1. A higher ARI value indicates a higher consistency between the clustering results and external labels.

$$NMI(X;Y) = \frac{I(X,Y)}{F(H(X),H(Y))}$$

$$I(X;Y) = \sum_x \sum_y p(x,y) \log \frac{p(x,y)}{p(x)\,p(y)}$$

NMI is a normalized form of Mutual Information, and NMI normalizes it to have a range between 0 and 1. Higher ARI and NMI values indicate a higher consistency between the clustering results and external labels. We calculate ARI and NMI using functions from the python package sklearn.metrics.

### Clustering internal indicators

For data without ground truth classification, the internal clustering metrics Calinski-Harabasz Index (CH), Davies-Bouldin Index (DB), and silhouette coefficient are used to evaluate the quality of clustering results.

CH index measures the compactness within clusters and separation between clusters. Higher CH values indicate better clustering results with tighter and more separated clusters. The silhouette coefficient is similar to CH. A higher value indicates better clustering results, with good separation and strong internal cohesion. While DB Index is the opposite, lower DB values indicate better clustering results. We calculate CH, DB and silhouette coefficient using functions from the python package sklearn.metrics.

**Spatial distribution indicator**

Local inverse Simpson's index (LISI) is a biodiversity metric that measures diversity and evenness within a specific area. It is the reciprocal of the Simpson's index. A higher value of the index indicates a more diverse and evenly distributed community, suggesting a higher level of spatial continuity. This index helps assess the spatial patterns of species distribution and identify areas important for maintaining ecological connectivity. We calculate LISI by writing a function using python (https://github.com/STOmics/StereoMMv1).

**Differential analysis and enrichment analysis**

We used fold changes and adjusted p values to identify differentially expressed genes (DEGs) according to the spatial domain (cluster label). For specific clusters, we compare gene expression within the cluster to gene expression across other clusters. By calculating the log2 fold change based on the average gene expression in these two groups, we can reveal the average expression differences. We calculate the aforementioned metrics using the FindMarkers function in Seurat package. For all the data, we define differentially expressed genes by setting thresholds of absolute value of log2 fold change (log2FC) $\geq 0.25$ and p value $< 0.01$.

Based on the results from the FindMarkers function, all genes are sorted in descending order based on their fold change. We then utilize the R package fgsea to perform gene set enrichment analysis (GSEA), which helps to identify gene sets that are significantly enriched in the differentially expressed genes. GSEA provides insights into the functional significance of the DEGs by examining their enrichment in predefined gene sets representing specific biological pathways or functions. In this case, the predetermined gene set is the HALLMARK gene set downloaded from the MSigDB website.

**Spatial colocalization analysis**

We applied Kernel Density Estimation (KDE) to determine the density distribution of point attributes within each spatial transcriptomics section. KDE is a statistical technique used to estimate the probability density function of a continuous random variable. It is a fundamental data smoothing problem where inferences about the population are made based on a finite data sample. For lung adenocarcinoma (LUAD) and lung squamous cell carcinoma (LUSC), we utilized either annotated or manually labelled categories. For marker genes, we estimated the density of points expressing these genes. The correlation coefficient was obtained by calculating the Pearson Correlation Coefficient (PCC) between the cell identity (LUAD or LUSC) and the corresponding marker gene's KDE. This PCC serves as a quantitative measure of the colocalization of the two factors within the tissue sections, reflecting how closely the cell types and marker gene expressions are associated in the spatial context.

The kernel function shapes the density estimate in KDE, influencing the smoothness and sensitivity of the analysis to the underlying data distribution.

The bandwidth parameter ‘h’ in kernel density estimation critically controls the smoothness of the density curve, balancing detail and generalization to avoid overfitting or excessive smoothing.

**WGCNA gene module analysis**

We describe the application of Weighted Gene Co-Expression Network Analysis (WGCNA) to identify clusters of highly correlated genes, known as modules, from large gene expression datasets. WGCNA identifies gene modules by constructing a weighted network from gene expression data through pairwise correlations and a scale-free topology. Once the network is constructed, genes are grouped into modules using average linkage hierarchical clustering coupled with the Topological Overlap Measure (TOM) to quantify network connectivity of genes. Modules are then related to external phenotypic traits by calculating the correlation between the module eigengene, which is the principal component representative of a module's gene expression profiles, and the trait of interest. This allows for the identification of modules that are potentially relevant to the trait, providing insights into the underlying biological processes and pathways.

**Functional and Pathway Enrichment Analysis**

We combined Over-Representation Analysis (ORA) and Gene Set Enrichment Analysis (GSEA) to delineate the functional roles and pathways associated with our genes. Using the clusterProfiler R package, we conducted ORA on a select gene list, adopting Gene Ontology (GO) terms that met the significance threshold of p-value < 0.05, thereby pinpointing biological processes with significant enrichment. Concurrently, we utilized the fgsea R package to assess the overall gene distribution within predefined gene sets, aiding in the revelation of broad gene expression patterns. For GSEA, we downloaded the HALLMARK gene sets from the Molecular Signatures Database (MsigDB). Pathways with a GSEA output of p-value < 0.05 were considered significantly enriched, with a Normalized Enrichment Score (NES) > 0 indicating an overall upregulation of the pathway, and an NES < 0 indicating an overall downregulation.

**Copy number variation (CNV) analysis and differentiation analysis**

The inferCNV R package was applied to detect Copy Number Variations (CNVs) within the study. This technique provides high-resolution identification of CNVs by contrasting the gene expression profiles of individual cells against a reference group of cells with an assumed normal karyotype. In this instance, points annotated as immune cells ("B-cells", "T-cells", "Myeloid", "Plasmablasts") were used as the reference cells for inferCNV inference. Upon obtaining the inferCNV output matrix (infercnv.observations.txt), the returned values were subtracted by one and then squared to calculate the CNV score for each point.

The CytoTRACE software package was employed to evaluate the differentiation status of cells within our dataset. A higher CytoTRACE score indicates a less differentiated state, suggesting that the cell has the potential to differentiate into multiple cell types.

**Cell type annotation**

Cell2location was used to spatially map cell types identified in single-cell RNA sequencing (scRNA-seq) data onto spatial transcriptome data. This approach requires a reference scRNA-seq dataset in which cell types have been previously identified and characterized. Here we download a single-cell atlas of human breast cancers[1] for reference and deconvolve the mixture of 9 immune cell types and non-immune cells in the Stereo-seq data with the Cell2location hyperparameter N_cells_per_location= 1, Detection_alpha=20. Assign the most abundant cell type to each cell.

**Survival analysis**

We utilized the Kaplan-Meier Plotter, an online tool that facilitates the assessment of the effect of genes on survival in various cancer types. The tool integrates gene expression data with survival information, allowing users to input specific gene symbols and select relevant patient cohorts for analysis. Kaplan-Meier survival curves are generated, and log-rank tests are performed to determine the statistical significance of differences in survival outcomes between patient groups stratified based on gene expression levels. For all cohorts, we stratified the groups based on the median expression of the genes, and log-rank p-values $< 0.05$ were considered to indicate statistically significant differences.

This PDF file includes: Figs S1 to S6

**Supplementary Figures**

Fig. S1. Comprehensive parameter testing and ablation studies.
Fig. S2. Spatial domains identified by each algorithm on the lung adenosquamous carcinoma dataset.
Fig. S3. Performance of each algorithm in identifying spatial domains on lung adenosquamous cell carcinoma dataset.
Fig. S4. Colocalization analysis of marker genes and spatial domains.
Fig. S5. WGCNA distinguishes gene modules in lung adenosquamous carcinoma.
Fig. S6. Domain 11 in breast cancer exhibits features of tertiary lymphoid structure.

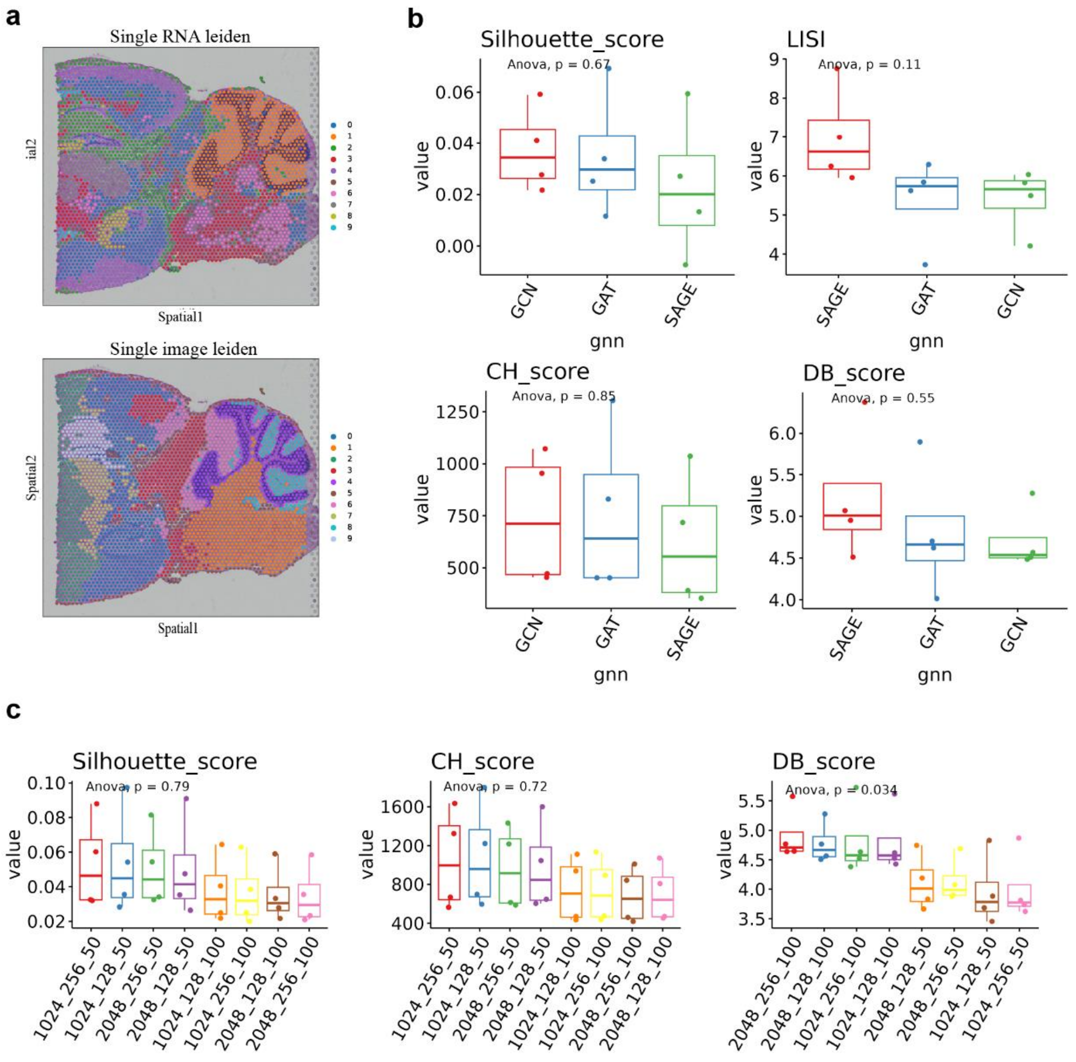


Fig. S1. Comprehensive parameter testing and ablation studies.

**a**, Spatial domain recognized by single modality, top: single transcriptome, below: single morphology. **b,** External clustering metrics and LISI scores for different types of GNNs. **c,** External clustering metrics with different numbers of nodes.

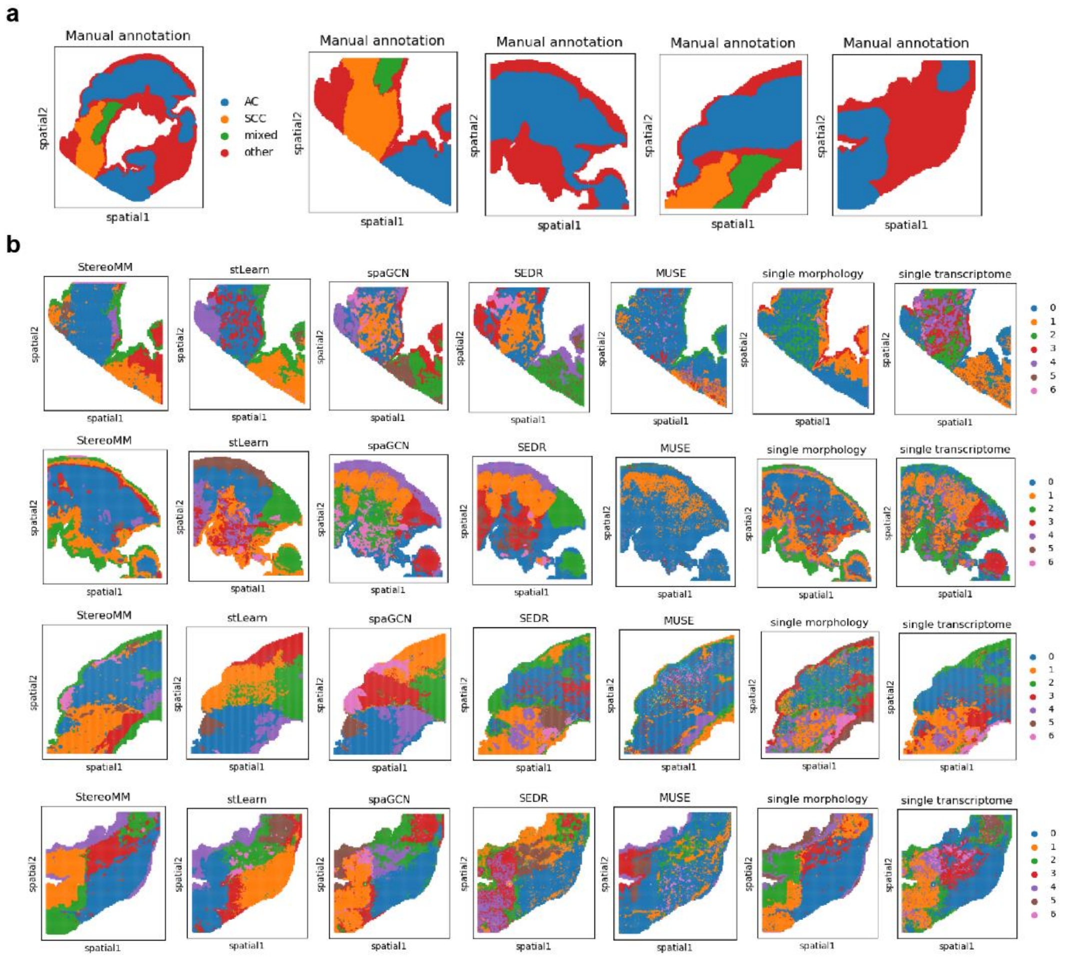


Fig. S2. Spatial domains identified by each algorithm on the lung adenosquamous carcinoma dataset.

**a,** The results of manual annotation. The left side is the display of the original single slide, and the right side is the display of the slide segmented into four slides. **b,** Spatial domains identified by all algorithms (StereoMM, stLearn, spaGCN, SEDR, MUSE, single morphology modality, single transcrioptome modality) for each slide.

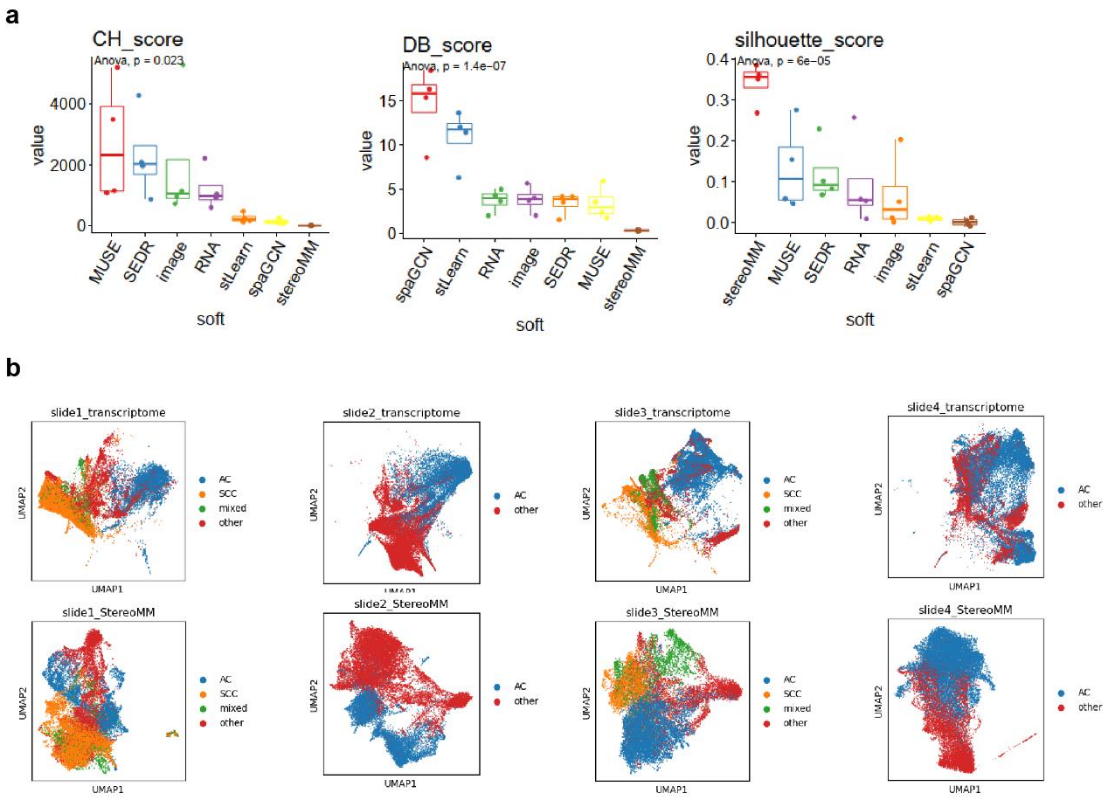


Fig. S3. Performance of each algorithm in identifying spatial domains on lung adenosquamous cell carcinoma dataset.

**a,** Boxplot of the cluster internal evaluation index for seven methods in all 4 slices. **b,** UMAP plots by raw transcriptome and StereoMM representation for each slide.

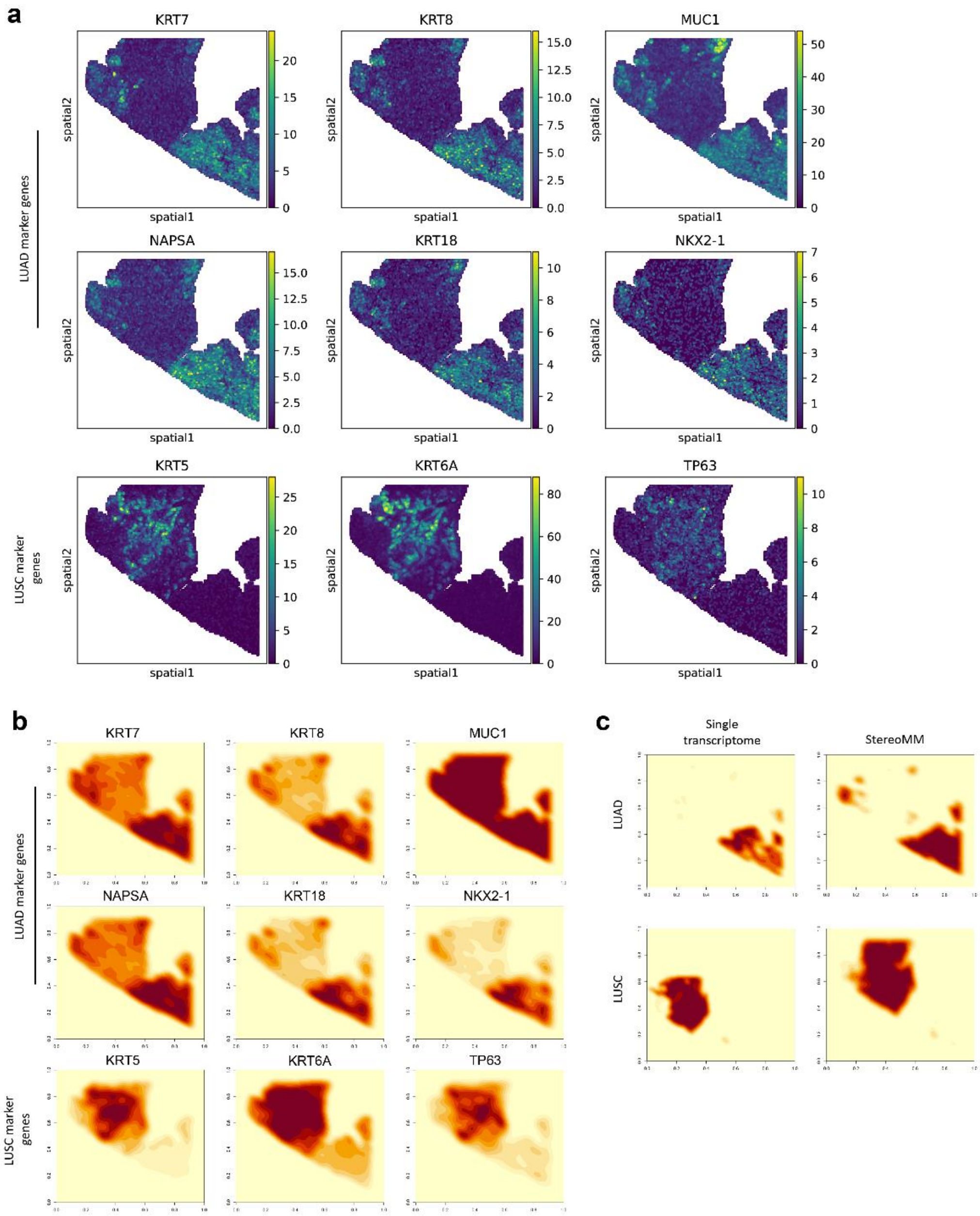


Fig. S4. Colocalization analysis of marker genes and spatial domains.

**a,** Spatial expression patterns of marker genes for LUAD and LUSC **b,** Kernel density maps for distribution statistics of LUAD and LUSC marker gene expression using the Kernel Density Estimation (KDE) method **c,** Kernel density maps were created using the KED method to delineate the spatial domains identified by stereoMM and individual transcriptomes in LUAD and LUSC.

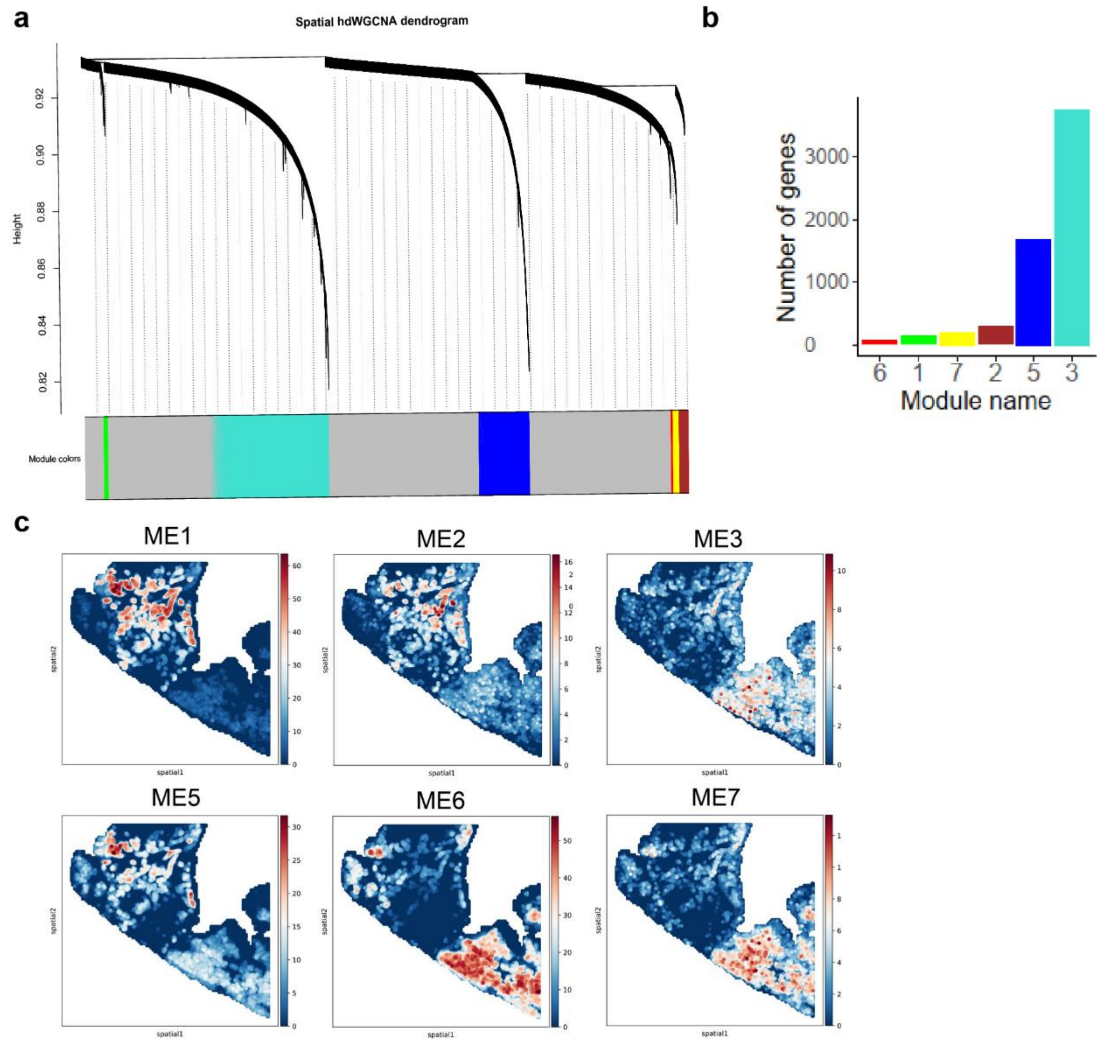


Fig. S5. WGCNA distinguishes gene modules in lung adenosquamous carcinoma.
**a,** WGCNA gene clustering diagram. The upper part is the hierarchical clustering dendrogram of genes, and the lower part is the gene module, which is the topological network module. **b,** Histogram of the number of genes within the WGCNA gene module. **c,** WGCNA gene module scores in situ.

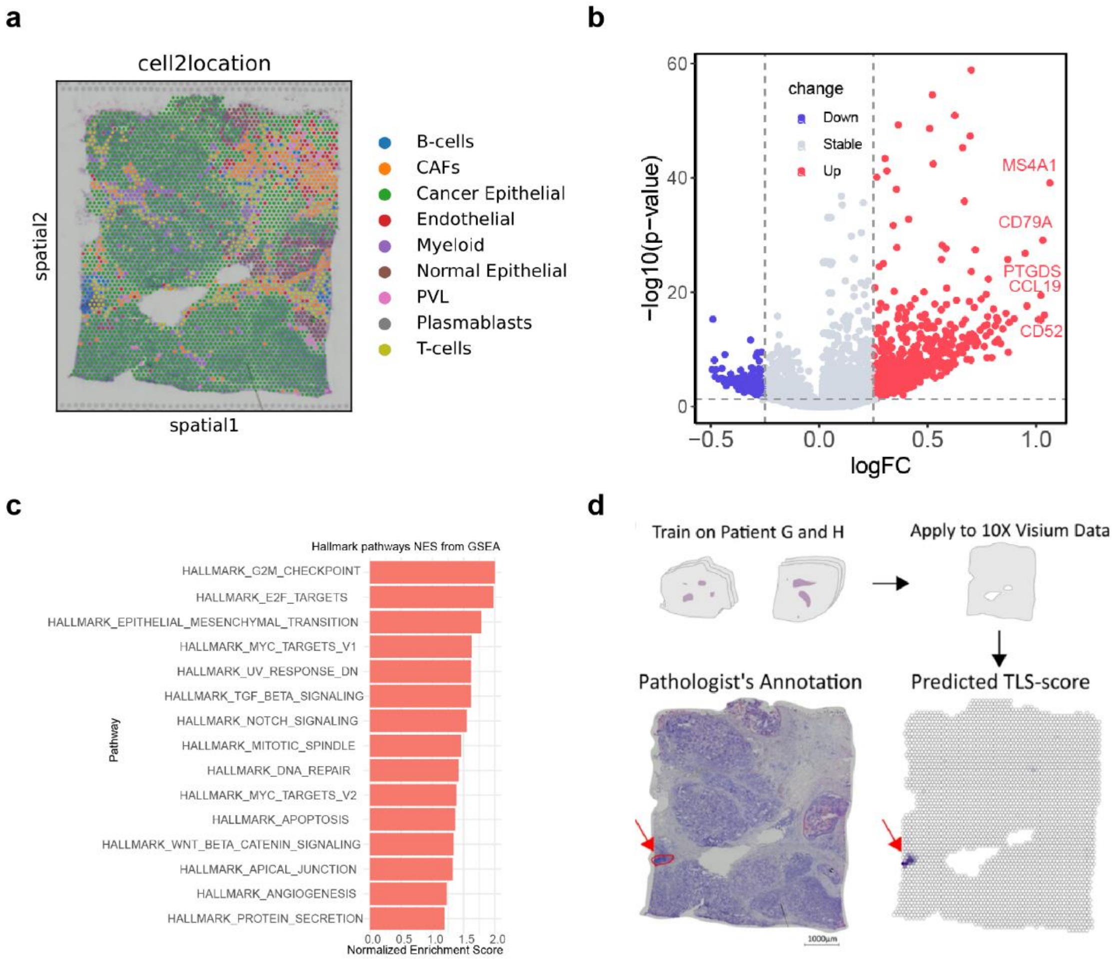


Fig. S6. Domain 11 in breast cancer exhibits features of tertiary lymphoid structure.

**a,** The spatial location of the cell2location annotation result. **b,** DEGs between domain11 and other domains. **c,** Gene Set Enrichment Analysis (GSEA) for fold change of genes expression. **d,** Area with high TLS scores reported in previous study.
